# Ecological stoichiometry and adult fat reserves suggest bet-hedging in *Drosophila melanogaster* development

**DOI:** 10.1101/780098

**Authors:** Tatjana Krama, Ronalds Krams, Priit Jõers, Māris Munkevics, Giedrius Trakimas, Severi Luoto, Sarah Eichler, David M. Butler, Enno Merivee, Anne Must, Markus J. Rantala, Jorge Contreras-Garduño, Indrikis Krams

## Abstract

The elemental composition of organisms relates to a suite of functional traits that change during development in response to environmental conditions. It may be a part of a phenomenon known as ‘developmental programming’, which hypothetically creates phenotypes that are better adapted to their environments. However, associations between developmental speed and elemental body composition are not well understood. We compared body mass, elemental body composition, food uptake and fat metabolism of *Drosophila melanogaster* Oregon-R male fruit flies across the time gradient of their larval development. The results showed that flies with intermediate and rapid developmental speeds were heavier than slowly developing flies. Slowly developing flies had higher body carbon concentration than rapidly developing and intermediate flies. Rapidly developing flies had the highest body nitrogen concentration, while slowly developing flies had higher body nitrogen levels than flies with intermediate speed of development. The carbon-to-nitrogen ratio was therefore lower in rapidly developing flies than in slow and intermediate flies. Feeding rates were lowest in the slowly developing flies. The amount of storage fats was highest in the intermediate group. This means that the growth of rapidly developing flies is not suppressed by stress and they actively convert the food they consume into growth with less emphasis on storage build-up, suggesting bet-hedging in the larval development. In contrast, flies in the intermediate developmental group had the greatest fat reserves which optimize fitness under many climatic conditions. Low food intake may slow down development and the accumulation of body fat reserves in slowly developing flies. However, at the cost of slower growth, their phenotype conceivably facilitates survival under higher stochasticity of their ephemeral environments spoiled by metabolic waste due to high density of conspecifics. Overall, this study suggests that bet-hedging may be a common developmental strategy in fruit flies to cope with environmental uncertainty.

## Introduction

Biotic and abiotic environmental stressors affect organismal development, with impacts that are able to create individual and population differences in life history strategies [1–4]. During stress, the internal state of an organism falls outside the organism’s typical operating range [5, 6]. Stress induces specific biochemical and physiological responses that counteract its consequences. Responses to adverse conditions represent cascades of internal changes to balance growth, reproduction, immune defense, self-maintenance and external stressors [7]. However, the availability of fitness-relevant resources in any particular environment is limited; time, effort and energy expended in any given way reduces their availability for other activities and processes [8, 9]. This often causes trade-offs in allocations of an individual’s resources to such competing life functions as immunity, reproduction, self-maintenance, development and growth especially under conditions of stress [10–12].

Ecological stoichiometry is a field of research that focuses on the interactions of organisms with their environments and the subsequent changes in the elemental composition of their bodies [9, 13, 14]. Organisms can regulate their internal state by adjusting food intake and metabolism, which have the potential to affect the elemental composition of the organisms [15]. Several environmental factors are known to influence elemental composition of organisms, including pollution [16, 17], ambient temperature changes [18, 19–21] and predation risk [20–23]. For example, predator-induced stress generally increases metabolic rate [7, 24–27]. Subsequently, rising energetic demands increase the overall demand for carbohydrate-based fuel and shift the metabolic balance away from anabolism that produce nitrogen-rich (N) proteins necessary for growth [7, 25]. Under such circumstances, the body utilizes proteins to produce glucose [7, 25, 28]. However, this is not universal condition: for example, *Drosophila melanogaster* fruit flies reared together with spiders had increased concentration of body N and lower body mass, while their body carbon (C) remained the same as in control individuals [22]. These stoichiometric changes improved climbing speed and adult survivorship under experimentally induced predation risk [22]. This may represent bet-hedging in anti-predator defense where fitness is higher under stress than in the typical environment.

It has been suggested that growth rates will always be at their physiological maximum unless there exists some intrinsic cost of rapid growth [29, 30]. However, evidence shows that few organisms develop at their maximum growth rate [31–33]. The pace-of-life syndrome (POLS) hypothesis [34–36] predicts that rapidly developing individuals with high activity have faster life histories (e.g., faster development and earlier reproduction) and higher metabolic rate, which increase resource acquisition [34] and reduce life span through increased oxidative damage [37]. Importantly, the growth rate hypothesis (GRH) predicts that rapidly growing organisms have higher concentrations of body N because of increased synthesis of N-rich proteins [13, 38]. On the other hand, rapid development can be adaptive because of bet-hedging, which promotes survival and reproduction under transient environmental changes [39, 40]. In bet-hedging, a single genotype produces an array of phenotypes: this risk-spreading strategy guarantees that some phenotypes optimally match the environment [41]. For example, it has been shown that an optimal strategy of the desert plant is to improve survival by hedging its bets and have seeds that start their growth as early as they can while the others germinate only under the most suitable conditions [42]. Bet-hedging may be especially useful in *D. melanogaster*, a species that has a relatively short generation time in relation to seasonal temperature fluctuations and the availability of resources. In *D. melanogaster*, another strategy to cope with environmental uncertainty is adaptive tracking where individuals survive changing conditions by having diversified phenotypes as a result of genetic variation. However, adaptive tracking has the potential to outcompete bet-hedging only under conditions of stable climate and predictable resource access [43].

Surprisingly, slow development may also incur costs. Slower development has been associated with upregulation of stress-related genes [44]. It has also been shown that wings of late-hatched female damselflies, *Lestes viridis*, are less symmetrical than those of early-hatched individuals suggesting developmental stress [45]. A recent study reported high N concentrations in the bodies of rapidly developing crickets [15], supporting the GRH [13, 38]. However, slowly developing crickets had significantly higher body C and higher C/N ratio (both are indicators of stress) than rapidly developing crickets, a finding that suggests higher stress levels in slowly growing individuals [15]. This may be because individuals that grow slower than conspecifics risk finishing their larval development in environments that are polluted with nitrogenous metabolic waste left by rapidly developing individuals. It has been shown that *D. melanogaster* individuals that eclose later than rapidly and intermediately growing ones have increased tolerance to urea/ammonia, which requires time and energy to develop specific biochemical adaptations [46, 47]. It therefore has the potential to increase physiological stress in slowly developing individuals.

In this study we investigated concentrations of body C, N and the C/N ratio in *D. melanogaster* larvae with slow, intermediate and rapid developmental speed. Developmental time is a trait of significant relevance to fitness especially in *Drosophila* species [48]. Fruit flies often occupy ephemeral habitats such as rotting fruits, which may promote rapid larval development as fruits dry out. However, selection for faster development and early reproduction leads to a reduction in egg-to-eclosion development time, survivorship, lower adult dry body mass at eclosion and a decline in larval growth rate [47]. Therefore, we did not use selective lines and did not manipulate the presumed causal factors of growth experimentally but took advantage of natural variation in studying the effects of elemental composition and body mass in male fruit flies across the time gradient of their larval development. We also investigated feeding rates and fat reserves of fruit flies from three different developmental groups. This was done because rapidly developing individuals are generally considered to consume more food to grow faster than slowly growing individuals [49]. Based on prior research [15], we predicted higher feeding rates, higher body N, lower C and lower C/N ratio in rapidly developing flies. With regard to bet-hedging in developmental speed [50], we predicted low fat reserves in rapidly and slowly developing flies because fat may be used to sustain high growth rates in the former individuals and to enable specific physiological survival mechanisms in deteriorated environments in the latter individuals [22]. We studied only males because a sizable fraction of the body of a mated female is made up of developing embryos, which have a significantly different metabolism than the female soma. Therefore, using females would lead to obfuscated results with regard to adult stoichiometry.

## Materials and methods

### Insects and development groups

Fruit flies were maintained in a lab at the University of Tennessee-Knoxville at 25 ± 1 °C under a constant 12:12 h light–dark cycle. The wild strains Oregon-R-modENCODE (#25211) of *D. melanogaster* were obtained from Bloomington Drosophila Stock Center (IN, U.S.A.).

The flies were isolated under carbon dioxide anaesthesia. To ensure virginity, we isolated females within 7 hours after imaginal eclosion. All flies used to produce progeny eclosed themselves on day 12. To obtain fruit flies for this study, we placed ten females and ten males per vial (n = 22; Flystuff Polystyrene vials: 24 inner diameter x 95 mm height) with 6 ml of food (cornmeal, dextrose and yeast medium) to copulate for 9-10 h. Anaesthesia was not used for this part of the study. We subsequently removed all females from vials and moved them to other vials of the same size and food volume (6 ml) for 1 hour to oviposit. Each vial contained food consisting of cornmeal, dextrose and yeast medium.

When the eggs began to eclose, we collected 10 flies of each of the following groups: 1) most rapidly developing flies, 2) most slowly developing flies and 3) flies that developed with intermediate speed. We collected a total of 30 flies per each vial. We used individuals with intermediate speed of development as controls to the rapid development and slow development groups. The most rapidly developing flies eclosed on day 9 (females laid eggs on day 0), the most slowly developing flies were collected on day 15, while we considered flies eclosing on day 12 as the intermediate group. Each vial contained between 73 and 80 larvae. This number is considered to represent low to average density of offspring for this vial size [51], suggesting the larvae did not experience severe competition for food and space as opposed to normal lab conditions where more than 150 larvae usually live in 24 x 95 mm vials. Removal of eclosing fruit flies affected the density of remaining larvae. To keep the density constant, we added a corresponding number of larvae and fresh food to each vial.

### Fruit fly body C and N content

Within 6-8 hours after imaginal eclosion, each fly was placed separately in vials for 1 hour with only water and no food provided. This ensured all consumed food and faeces were released during the fasting period. Then the flies were frozen at −80 °C. On the next day, the flies were dried at 75 °C for 72 hours, and weighed as groups of 10 males [52] using a Sartorius MC5 microanalytical balance with an accuracy of ±1μg. Dry body weight was calculated for individual flies as the dry mass of each replicate divided by the number of flies assigned in each replicate. Each group of 10 individuals was ground to a homogenous powder.

The percentage of C and N content was measured from the mass of whole flies using a C/N combustion auto-analyser [7, 22, 25]. Samples of C and N concentrations were measured as groups of 10 fruit flies equally representing each vial. In total, we measured 220 males for each developmental speed, with a total of 660 flies.

### Food uptake

We used additional six vials to collect fruit flies for feeding experiments. For each experiment, 150 flies were distributed to six separate vials (25 flies per vial) with standard food. The flies were allowed to recover overnight from CO_2_ exposure. On the next day we transferred flies to new vials without using gas. We transferred half of the flies to vials with standard food, whereas the other half was transferred to vials with food supplemented with 1% Blue FCF dye (Acros). After 1.5 h, 20 flies were collected and lysed by grinding in a mortar and pestle in 800 μl of phosphate-buffered saline (PBS). Debris was pelleted at 10,000 g_max_ for 10 min at 4 °C and supernatant was transferred to fresh eppendorf tubes without disturbing the pellet. Supernatant was centrifuged again as above, after which 400 μl of each supernatant were transferred to 2 wells (200 μl each) of 96-well plates. Absorbance of lysate was measured at 650 nm with Tecan Infinite M200 Pro spectrophotometer and amount of Blue FCF ingested was calculated using serial dilution of Blue FCF as a calibrator. There were seven vials for the rapid development group, five vials for the intermediate group and six vials for the slow development group.

### Triglyceride measurements

Ten flies were homogenized inside an eppendorf in 400 μl of PBS and 0.1% Tween 20. Lysate was heat-treated at 70 °C for 5 minutes to denature proteins after which one 20 μl aliquot was mixed with 20 μl Triglyceride Reagent (Sigma T2449) and one 20 μl aliquot was mixed with 20 μl of PBS and incubated at 37 °C for 30 minutes. During the incubation time, protein concentration in lysate was determined with Bradford assay. After incubation, samples were centrifuged for 15 sec at 14,000 g_max_ and 30 μl was transferred to a 96-well plate. PBS-treated samples were mixed either with 100 μl of Free Glycerol Reagent (Sigma F6428, determination of free glycerol) or water (background) and Triglyceride Reagent-treated samples were mixed with 100 μl of Free Glycerol Reagent (determination of total glycerol). Plates with samples were incubated at 37 °C for 5 minutes and the amount of quinonemine dye was measured with a spectrophotometer at 540 nm.

### Statistics

We used one-way analyses of variance (ANOVAs) to compare individual body mass, elemental and triglyceride composition, and food uptake in fruit flies. We used developmental group (rapid, intermediate, slow) as a fixed factor. We report only main effects if we did not find a significant difference between developmental groups; otherwise, we also report Tukey HSDs. Analyses were performed using R, version 3.5.2 [53]. Differences were considered statistically significant at *P* < 0.05.

## Results

### Body mass

Body mass of males in the three developmental groups was heterogeneous (one-way ANOVA: *F*_2,63_ = 60.08 *P* < 0.0001). While rapidly developing males (0.23 ± 0.015 g; mean ± SD) did not differ (Tukey HSD: *P* = 0.425) in their dry body mass from males in the intermediate development group (0.222 ± 0.03 g; mean ± SD), males in these both groups were significantly heavier (Tukey HSD: both *P*s < 0.0001) than males in the slow development group (0.166 ± 0.014 g; mean ± SD) (Fig. 1).

**Fig. 1.**
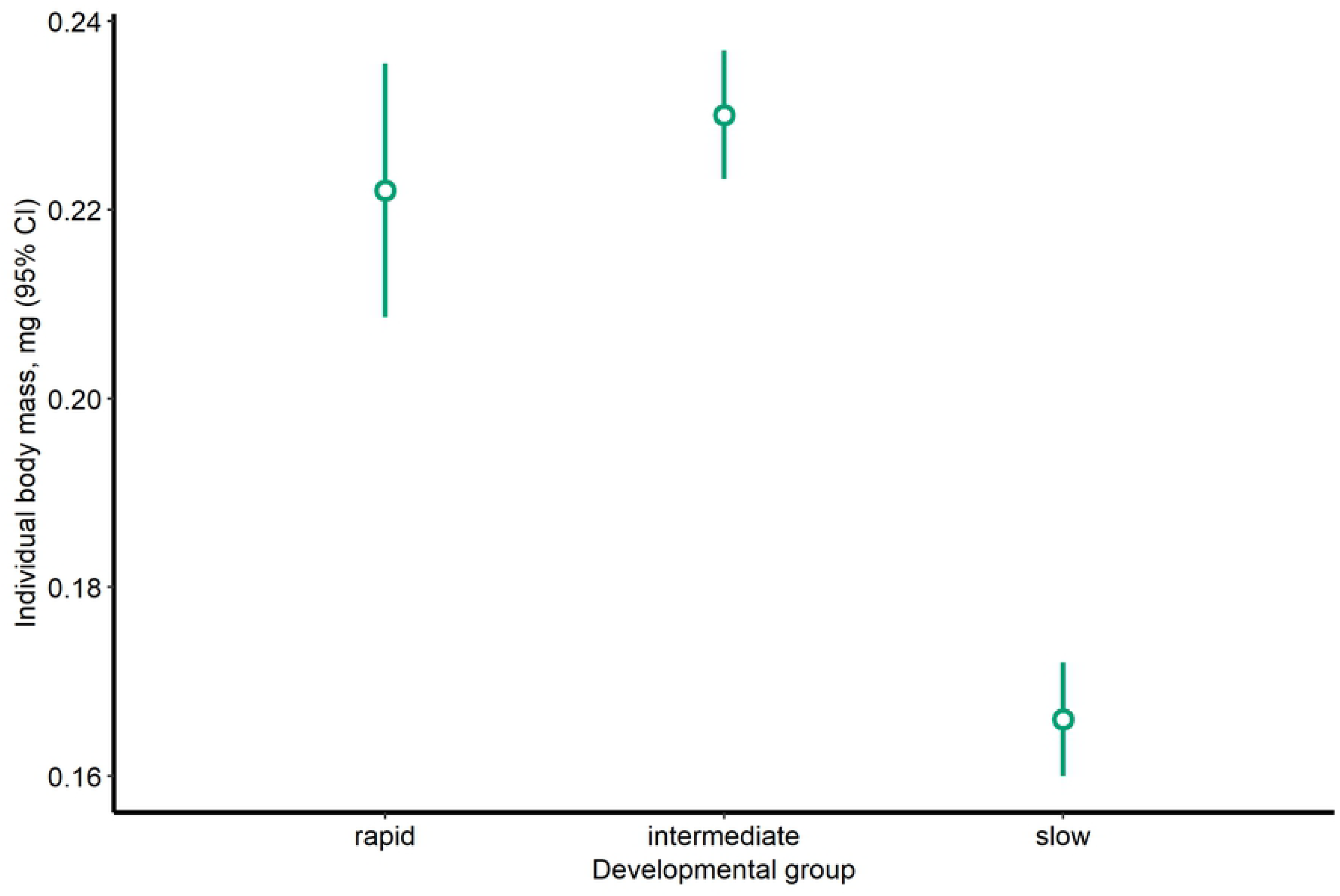
The average dry body mass of male fruit flies. Error bars represent 95% confidence intervals.

### Food uptake

The developmental groups significantly differed in their food uptake (one-way ANOVA: *F_2,15_* = 19.9, *P* < 0.0001). Although the food uptake of rapidly developing flies (0.33 ± 0.029; mean ± SD) did not differ significantly from the intermediate group (0.349 ± 0.069; mean ± SD) (Tukey HSD: *P* = 0.74), flies with rapid and intermediate development significantly differed from slowly developing flies (0.202 ± 0.030; mean ± SD) (Tukey HSD: both *P*s < 0.001) (Fig. 2).

**Fig 2.**
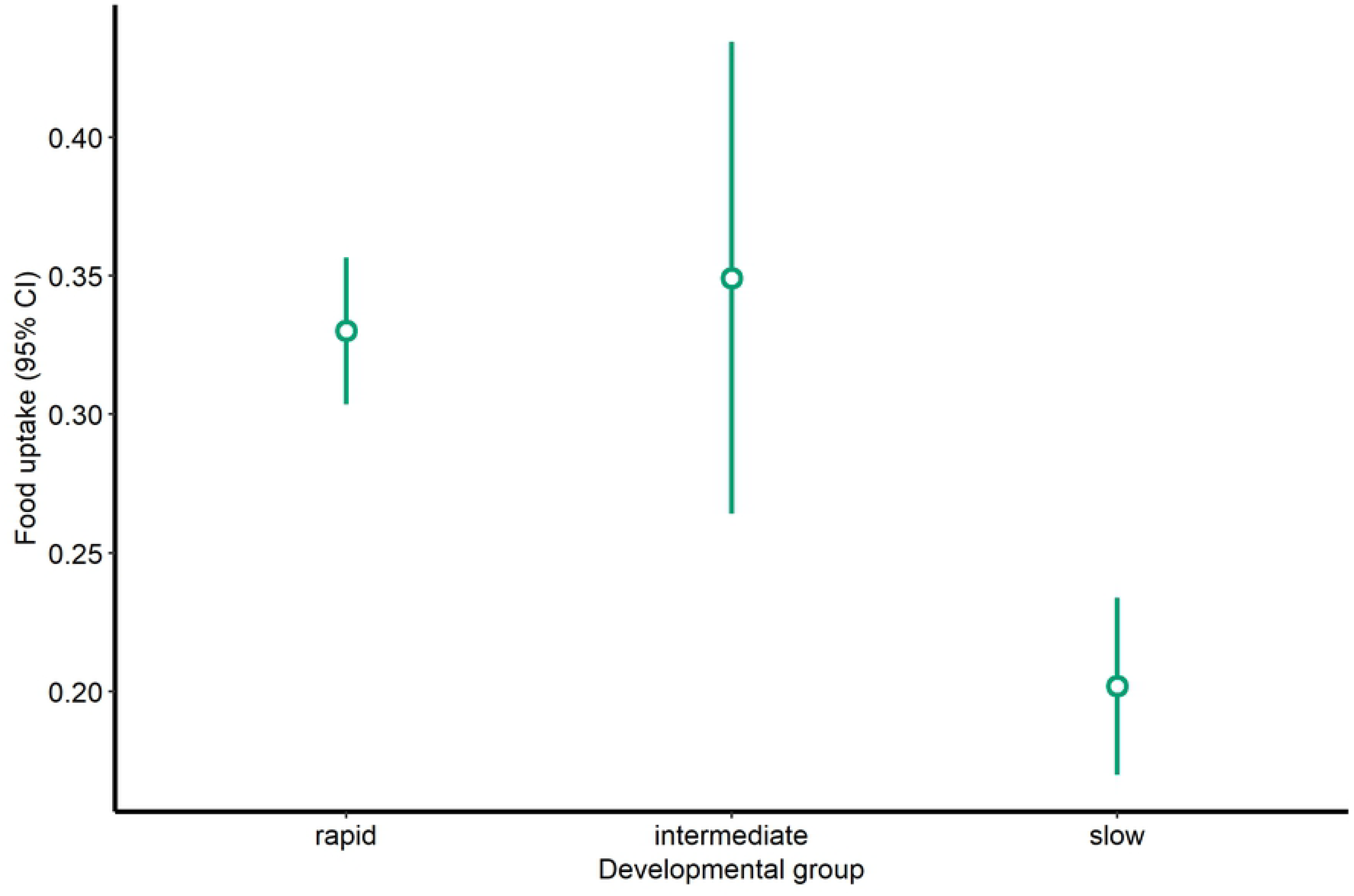
Feeding rates of fruit flies in rapid, intermediate and slow developmental groups. Error bars represent 95% confidence intervals.

### Carbon

There was a statistically significant difference between developmental groups in body C (one-way ANOVA: *F*_2,63_ = 25.77, *P* < 0.0001). Slowly developing fruit flies had higher body C (%) (54.698 ± 1.203, mean ± SD) than rapidly developing (52.661 ± 1.132) and intermediate (52.761 ± 0.807) fruit flies (Tukey HSD: all *P*s < 0.0001). Fruit flies in the rapid and intermediate developmental groups did not differ in body C (*P* = 0.948) (Fig. 3).

**Fig 3.**
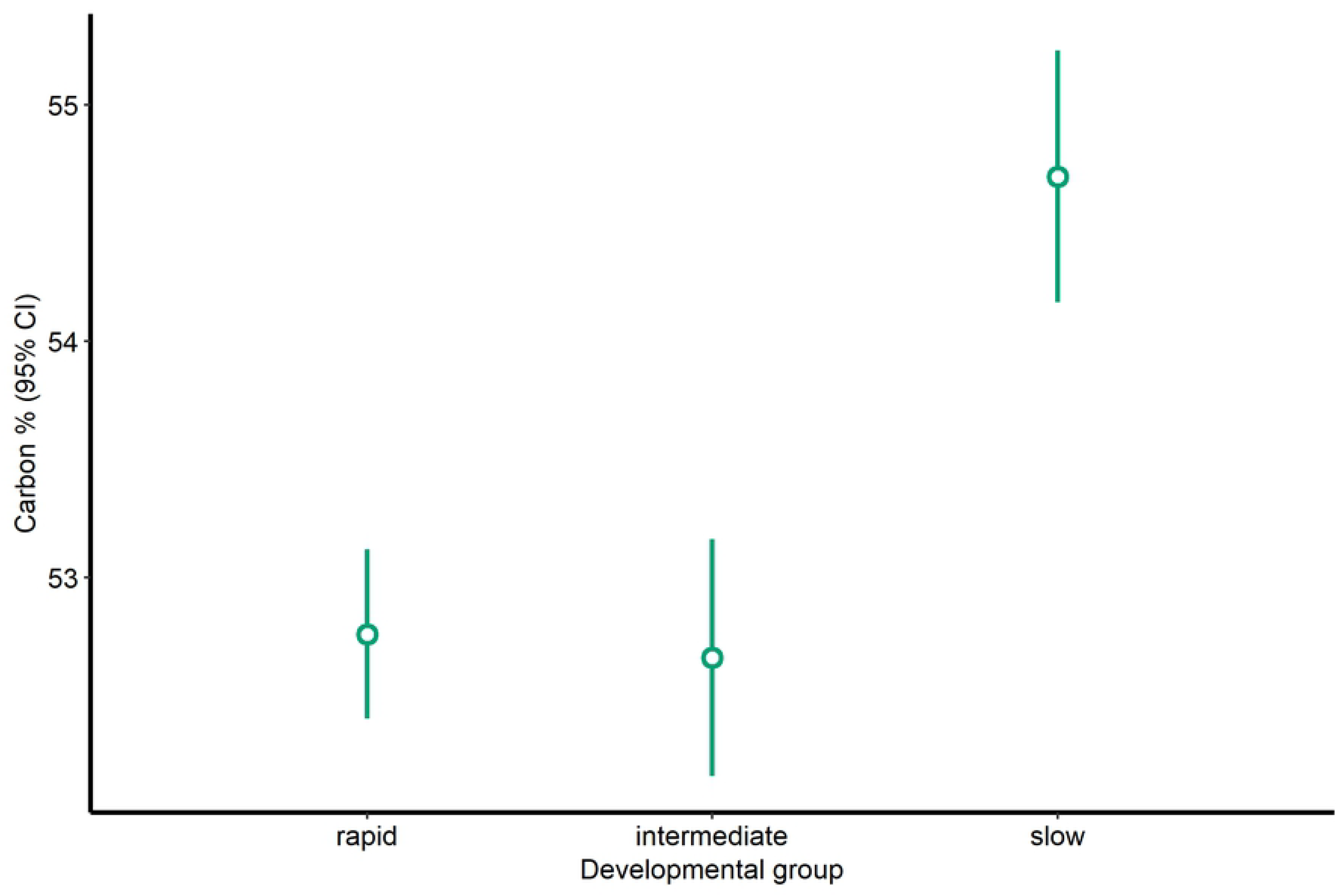
Average carbon percentage in adult *D. melanogaster* of slow development (*n* = 22), rapid development (*n* = 22) and intermediate development (*n* = 22) groups. Error bars represent 95% confidence intervals.

### Nitrogen

The effect of developmental group to body N was significant (one-way ANOVA: *F*_2,63_ = 92.87, *P* < 0.0001). Rapidly developing flies had higher body N (%) (10.239 ± 0.485, mean ± SD) than slowly developing (9.569 ± 0.360) and intermediate (7.769 ± 0.891) flies (Tukey HSD: *P* < 0.01 and *P* < 0.0001, respectively). Fruit flies in the slow and intermediate developmental groups differed in their body N (Tukey HSD: *P* < 0.0001) (Fig. 4).

**Fig 4.**
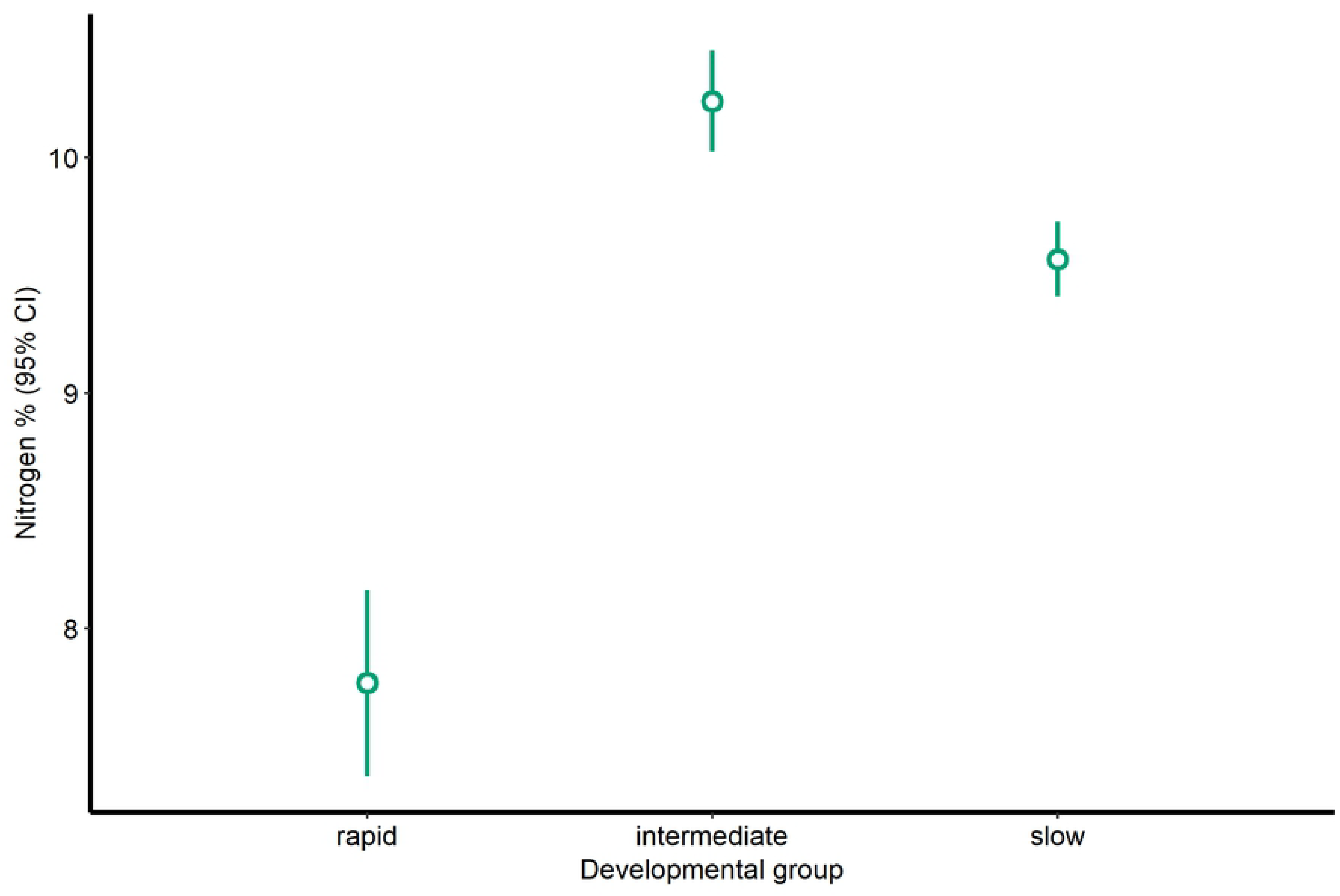
Average nitrogen percentage in adult *D. melanogaster* of slow development (*n* = 22), rapid development (*n* = 22) and intermediate development (*n* = 22) groups. Error bars represent 95% confidence intervals.

### C/N ratio

The effect of developmental group was significant (one-way ANOVA: *F*_2,63_ = 80.71, *P* < 0.0001). Rapidly developing fruit flies had lower C/N ratio (5.152 ± 0.224, mean ± SD) than slowly developing (5.726 ± 0.282) and intermediate (6.867 ± 0.703) fruit flies (Tukey HSD: *P* < 0.001 and *P* < 0.0001, respectively). Fruit flies in the slow and intermediate developmental groups also differed in their C/N ratio (*P* < 0.0001) (Fig. 5).

**Fig 5.**
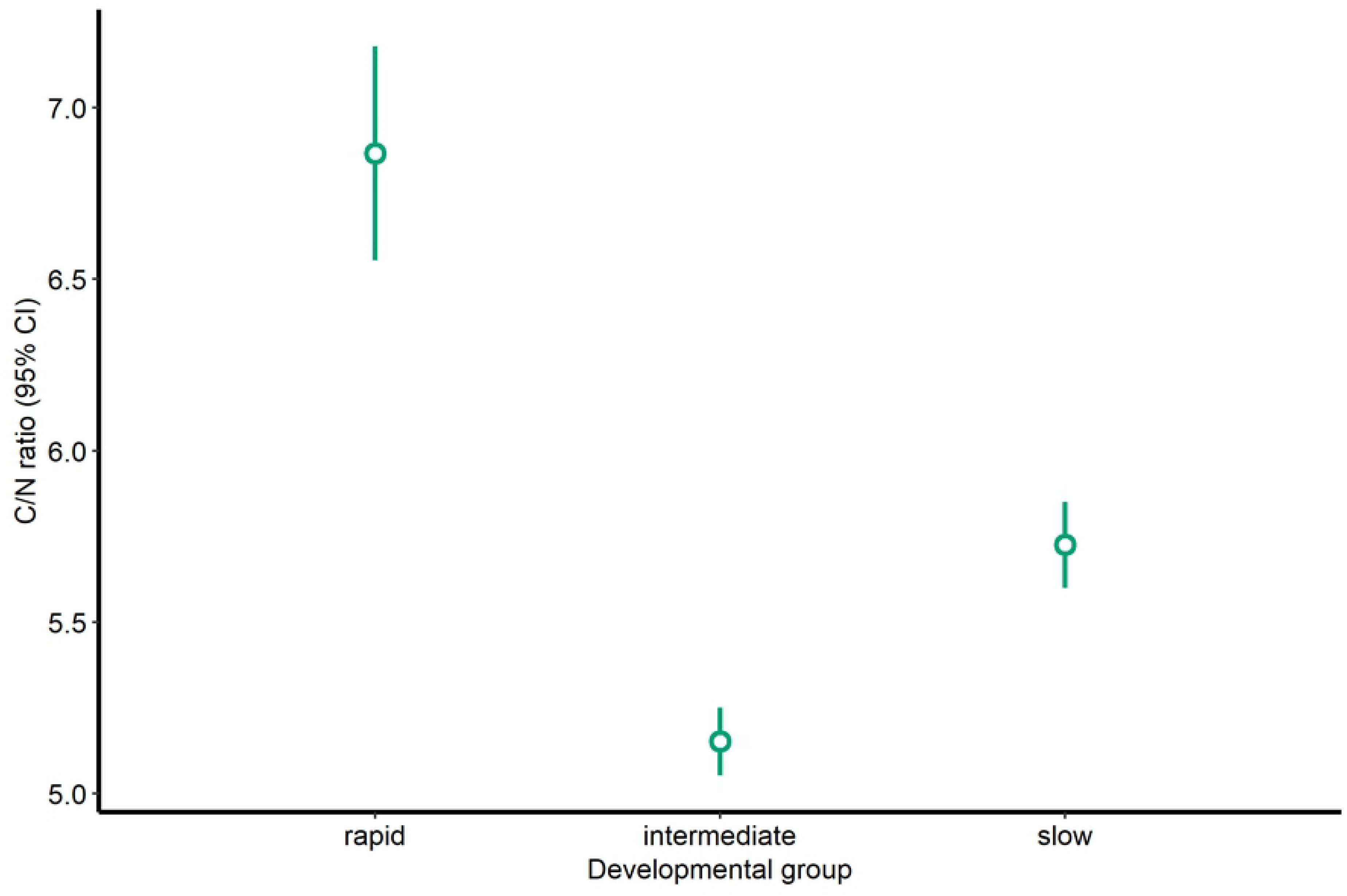
Average carbon-to-nitrogen ratio (C/N ratio) in adult *D. melanogaster* of slow development (*n* = 22), rapid development (*n* = 22) and intermediate development (*n* = 22) groups. Error bars represent 95% confidence intervals.

### Fat metabolism

The effect of developmental group on triglycerides was significant (one-way ANOVA: *F*_2,12_ = 35.5, *P* < 0.0001). Triglycerides (μg glycerol/mg protein) in rapidly (16.772 ± 1.127, mean ± SD) and slowly (18.609 ± 1.178, mean ± SD) developing flies were lower than in intermediate (42.892 ± 9.335, mean ± SD) flies (Tukey HSD: both *P*s < 0.0001; Fig. 6 A). Triglycerides did not differ in rapidly and slowly developing flies (Tukey HSD: *P* = 0.858; Fig. 6 A).

**Fig 6.**
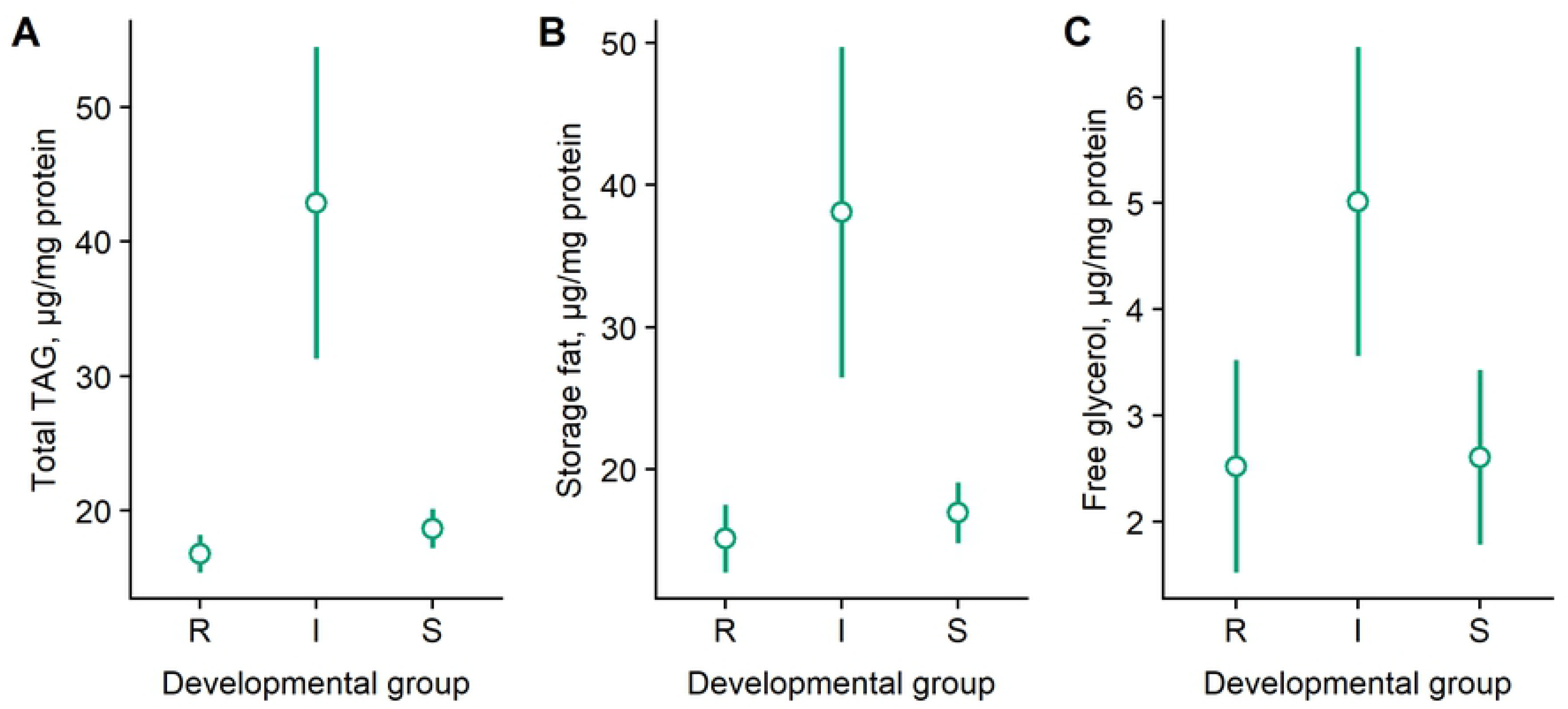
Triglycerides, storage fat and free glycerol in rapid (R), intermediate (I) and slow (S) development fruit flies. Error bars represent 95% confidence intervals.

The effect of developmental group on storage fats was significant (one-way ANOVA: *F*_2,12_ = 25.93, *P* < 0.0001). Storage fats (μg glycerol/mg protein) in both rapidly (15.149 ± 1.915, mean ± SD) and slowly developing flies (16.953 ± 1.705, mean ± SD) were lower than in the intermediate group (38.080 ± 9.353, mean ± SD) flies (Tukey HSD: *P* < 0.0001 and *P* < 0.001, respectively; Fig. 6 B). Rapidly and slowly developing fruit flies did not differ from each other (Tukey HSD: *P* = 0.868; Fig. 6 B).

The effect of developmental group on free glycerol in fruit flies was significant (one-way ANOVA: *F*_2,12_ = 12.2, *P* < 0.01). Free glycerol (μg glycerol/mg protein) was also lower in both rapidly (2.520 ± 0.806, mean ± SD) and slowly (2.605 ± 0.663, mean ± SD) developing flies (Tukey HSD: both *P*s <0.01; Fig. 6 C) compared to flies in the intermediate developmental group (5.014 ± 1.173, mean ± SD). Rapidly and slowly developing flies did not differ in free glycerol (Tukey HSD: *P* = 0.988; Fig. 6 C).

## Discussion

The whole life cycle of *D. melanogaster* from egg to adult is variable and normally takes between 9-15 days. Most fruit flies eclose around day 12 at 25 °C. Our results show that this considerable variation in the developmental time of fruit flies is associated with substantial between-individual variation in food uptake, body mass and the elemental composition and fat reserves of their bodies. The findings reported here are consistent with a recent study on the effects of developmental speed on the stoichiometry of crickets (*Gryllus integer*) [15]. Because of the high variability of eclosion time, low larval density and other conditions kept relatively constant, our results suggest that developmental plasticity and adaptive tracking alone cannot explain the observed variability of phenotypes. We suggest that the observed diversity of phenotypes was produced by developmental stochasticity where a single genotype generates an array of adult phenotypes regardless of variation in environmental conditions [11, 41]. This does not exclude the possibility that the observed variation in developmental time is a result of a combination of plasticity, adaptive tracking and bet-hedging [54]. However, it is important to note that all flies used in this study were progeny of adults that had intermediate developmental speed. Furthermore, the differences in elemental composition, food uptake, fat reserves and body mass were observed in fruit flies grown in low densities in identical laboratory conditions. Under phenotypic plasticity, the flies would be expected to follow similar developmental patterns as a response to similar environments, which was not the case.

Although rapidly growing fruit flies developed for the shortest time, these flies were heavier than slowly developing ones. The high developmental speed is explained by the high feeding rates and low storage fats of rapidly developing flies [49]. Importantly, high body N, low C and low C/N ratio suggest that rapidly developing fruit flies experienced low levels of stress, a finding that appears to contradict results from a number of studies which show that rapid growth is costly [33, 55–57]. Our results suggest that growth rates in rapidly growing *D. melanogaster* are optimized rather than maximized and that these rates most likely do not operate near their physiological maximum [8, 10, 58] where rapid growth cause oxidative stress and other costs [24, 33, 59]. This idea is further supported by Prasad and co-authors [47] showing that rapid development in selective lines is associated with low adult body mass only upon selection.

Specifically, we show that rapid and slow development represents bet-hedging in fruit flies whose parents eclosed on day 12. Intermediate development might be considered as adaptive tracking where individuals survive changing conditions by having diversified phenotypes as a result of genetic variation. Although flies of the intermediate group eclosed on day 12, we cannot exclude adaptive tracking in these flies because (genetically similar) isofemale lines were not used in this study. The low fat reserves in rapidly developing flies may indicate that these individuals have invested their nutritional reserves to eclose early or, alternatively, that they have never had any reserves to deposit. Early eclosion is a valuable strategy when there is high uncertainty in environmental conditions and fitness can be achieved if individuals develop and reproduce as fast as possible [43]. Moreover, larger bodies in insects usually indicate higher reproductive potential [60] or better immunity [61–63], both of which may be crucial in the case of unpredictable climate conditions and/or ephemeral food resources [64]. In contrast, adaptive tracking is suitable for individuals that eclose later in the season when consistent warm conditions are more likely than in the beginning of breeding season or in low latitudes [65]. Individuals following the adaptive tracking strategy may make trade-offs between their body size and body energy reserves to achieve the highest fitness; they can also afford the state of dormancy to survive unfavorable environmental conditions [66], which is less likely in individuals with much lower internal energy reserves.

Slowly developing fruit flies had significantly higher body C, lower N and higher C/N ratio than rapidly developing flies. This finding suggests that slowly developing flies are under higher stress levels during their larval development than rapidly developing flies. Stress is supposed to increase metabolic rate [15, 59; 67] and energetic demands by raising the need for carbohydrate fuel. This increases body C content and lowers N because stress responses require the breakdown of body proteins to produce C-rich glucose [7, 25, 28]. However, our results do not fully support previous findings of stress ecology [68–70] because rapidly developing flies had low body N and low body C. Importantly, the feeding rate of slowly developing males was the lowest of all three groups, while their body N was significantly higher than that of flies in the intermediate developmental group. The C/N ratio of slowly developing flies was lower than that in the intermediate developmental group, and their body C was higher than in the intermediate group. Our results indicate that the high body N lowered the C/N ratio of slowly developing flies compared to the intermediate developmental group. Although it is possible to propose that high C/N ratio in slowly developing flies was caused by low fat reserves, this does not explain the high body C concentrations in these flies. Overall, this shows that the observed simultaneous increase in C and N may render the C/N ratio a less reliable stress indicator than in such cases where the concentration of only one element substantially changes in response to environmental changes.

It is important to understand possible reasons for the heightened N content in slowly developing flies given their low food uptake, small sizes and high body C. It has been shown that slow development usually results in deteriorated larval environment because of higher conspecific density, the presence of larger rapidly developing larvae and relatively higher levels of metabolic waste left by more rapidly developing larvae and slow developing flies themselves [46, 71]. Slowly developing flies need to create specific biochemical/physiological adaptations against increased toxicity in their environment. Previous work has demonstrated that populations reared under crowded larval conditions evolve increased resistance to both urea and ammonia [72, 73], which has the potential to affect body composition of slowly developing flies. However, flies need selection to reach increased resistance to metabolic waste [49]. It is not clear whether the relatively low larval density in our study was enough to impair food quality: we therefore cannot ascertain that slower development was induced as an insurance against an unavoidable increase of urea/ammonia in the food/environment of slowly developing flies in our study.

Urea and ammonia have cytotoxic effects on *D. melanogaster*. Urea is a protein denaturant [74] and the larvae reared on urea-containing media have increased levels of proteins containing isoaspartyl residues which itself can be responsible for the increased concentration of body N in slowly growing flies [75]. Ammonia is known to have neurotoxic properties [76]. It accumulates to significant levels in larval cultures [46], which makes ammonia a potential candidate for the observed high levels of body N in slowly developing flies. Moreover, the expression of specific adaptations to develop and live in toxic environments requires the production of protein cosolutes that act as cytoprotectants [77]. These substances may maintain high body N in slowly developing flies via their protein stabilizing effects [78]. Future studies on fruit fly adaptations have to focus not only on protein presence but also on proteostasis. Finally, urea may be used as an important N source and it can be assimilated through symbiotic interactions with a variety of ureolytic microorganisms [79]. Thus, bacteria harvesting intestinal urea may also be responsible for the accumulation of N and the heightened body N in slowly developing flies.

The conditions experienced during larval growth and metamorphosis are important because neural and other physiological systems are under development and can be shaped by different types of stress [80]. The expression of bet-hedging and the developmental channeling of organisms into rapid or slow growth may cause permanent changes and lead to functionally coordinated sets in a range of phenotypic traits that benefit from the synergistic coordination of such traits [cf. 11, 81, 82]. In this study, we show that differences in developmental speed in fruit flies result in differences in body mass, C and N content and C/N ratio, food uptake, body mass and the amount of fat reserves. This knowledge may contribute to research on ‘developmental programming’, since variation in developmental speed can be a mechanism that creates phenotypes which are maximized for fitness [15, 80, 83, 84]. Our results also highlight the need to assess the concentration of body phosphorus to better understand the observed variations of C and N in *D. melanogaster*: phosphorus is important for RNA production and serves (quantified as the RNA:DNA ratio) as a proxy for protein synthesis [17] and other life-supporting processes such as memory [85].

## Conclusion

It has been traditionally considered that fruit flies are subject to directional selection for rapid development under natural conditions [49, 86]. We did not observe that any noticeable decrease in body mass characterized rapidly developing fruit flies compared with the intermediate group. The elemental composition of rapidly developing flies showed no signs of elevated stress. This suggests that developmental speed may be a function of stabilizing rather than directional selection in fruit flies. Rapid and slow growth can be considered as bet-hedging strategies: rapid development facilitates coping with climatic unpredictability and allows reproduction during short periods of suitable weather, while slow development may optimize fruit fly phenotypes for an environment deteriorated by metabolic waste products. Based on our results, we predict a substantially narrower variation in developmental speed, the amount of body reserves, elemental composition and behavior [65] in fruit flies that live in more predictable and warmer climates. Taken together, our findings suggest that developmental speed is an important variable influencing elemental composition and that both developmental speed and stoichiometry belong to a broader suite of life history traits [cf. 11, 87, 88]. Overall, this study indicates that a combination of biochemical, physiological and behavioral approaches [15, 81, 82] is required to further develop the general stress paradigm, while focusing on ecological stoichiometry can provide an improved understanding of the dynamic ways in which organisms adapt to evolutionary selection pressures.

## Acknowledgements

This work was supported by Fulbright Program of the Department of State of the US, Latvian Science Council (lzp-2018/1-0393) and a personal grant (PUT1223) from the Estonian Ministry of Education and Science.

